# Robust Feature Selection strategy detects a panel of microRNAs as putative diagnostic biomarkers in Breast Cancer

**DOI:** 10.1101/2023.07.03.547377

**Authors:** Maria Claudia Costa, Teresa Maria Rosaria Noviello, Michele Ceccarelli, Luigi Cerulo

## Abstract

MicroRNAs represent a comprehensive class of short, single-stranded, non-coding RNA transcripts able to interfere with the translation of their targets or degrade them. Since these molecules are dysregulated in several malignancies, they represent reliable biomarkers in various contexts. In this study, the application of several Feature Selection methods uncovers a panel of 20 microRNAs, of which *hsa-mir-337, hsa-mir-378c, and hsa-mir-483* are still poorly investigated in the context of Breast Cancer. This signature is capable of discriminating between healthy and tumoral samples, showing better classification performance when compared with differentially expressed microRNAs. Furthermore, a network-based centrality analysis on the gene targets of these transcripts highlighted *CDC25, TPX2, KIF18B, CDCA3, TGFBR2, CAV1, TNS1*, and *LHFPL6* as key dysregulated genes. This study provides *in silico* hypotheses as new insights for future *in vivo* or *in vitro* studies uncovering the role of these putative diagnostic biomarkers in Breast Cancer.

## 1 Introduction

Nowadays, the search for new, effective, and reliable diagnostic biomarkers is constantly growing, accompanied by the development of innovative computational techniques that can help their identification. Among the potential candidates useful for this purpose, non-coding transcripts are actually assuming an ever-increasing importance in this regard, particularly MicroRNAs. In addition to being pivotal in regulating a plethora of biological aspects, alterations in the expression of these molecules have been associated with several types of cancer [1]. Differential Expression Analysis represents a valid and effective method for the detection of key biomarkers in various pathological contexts, nevertheless, the use of machine learning-based Feature Selection (FS) strategies allows extracting panels of diagnostic molecules in an ever more efficient way, revealing key features often underestimated by canonical biological techniques. In this study, we analyzed three Feature Selection approaches in terms of stability and classification performance to detect a robust diagnostic panel of microRNAs in a Breast Invasive Carcinoma cohort. The Support Vector Machine - Recursive Feature Elimination (SVM-RFE) method returned the most robust signature, showing the highest classification performances, also when compared to the top differentially expressed (DE) microRNAs. Further, we applied various bioinformatics techniques in order to have more insights into the biological contexts in which these microRNAs act: once the microRNAs-gene targets have been collected, a network-based centrality analysis allowed us to define the putative hub genes dysregulated by these key molecules, conceivably crucial for the disease to spread and progress. The complete pipeline is summarised in Figure 1.

**Figure 1:**
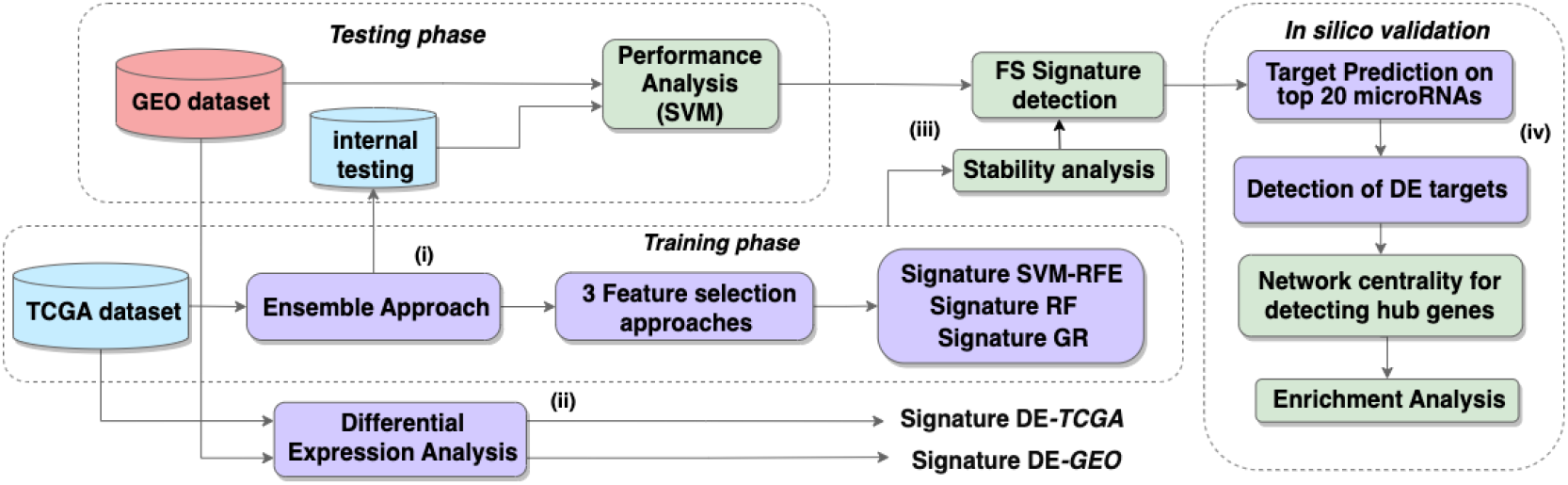
Pipeline overview. The workflow develops on four key points: Ensemble-FS computation on the training TCGA sub-datasets (i); DE analysis on the entire TCGA/GEO datasets (ii), classification performance analyses comparing DE and FS results and stability evaluation of the FS methods (iii); in silico validation of the top-20 microRNAs of the chosen signature, to detect the hub gene targets (iv).

## 2 Data and Methods

We collected 1881 microRNA-Seq data counts from the GDC data portal, TCGA project 1, Breast Invasive Carcinoma (BRCA) cohort, related to 300 solid primary tumoral samples (T) and 101 Normal Adjacent to the Tumor (NAT) samples, both belonging to ductal and lobular breast tissues; then a variance stabilizing normalization was performed on data prior the FS application. An independent validation dataset of 2565 microRNAs was downloaded from the Gene Expression Omnibus (GEO) 2 data repository (GSE97811), a microarray dataset, comprising 16 normal and 45 tumoral samples, then subjected to data imputation. Since this dataset refers to mature microRNA expression, in contrast to the TCGA data that includes precursor forms, we only selected the alternative mature microRNAs whose mean counts value across all the samples resulted higher than its opposite strand; also, microRNA names have been converted to corresponding precursor ones. This procedure reduced the dimensionality of the dataset to 1361 microRNAs. Corresponding TCGA RNA-Seq data was also collected and comprised a total of 20404 genes.

### 2.1 Feature Selection approaches and Ensemble procedure application

With the aim of comparing Filter-based, Embedded and Wrapper Feature Selection strategies, we chose Gain Ratio [2], Random Forest [3] and SVM-RFE [4], respectively belonging to each category, to be applied on 500 sub-datasets of the microRNA-Seq expression TCGA data in order to detect a robust panel of features discriminating with best performances between the Normal and the Tumor samples. We used 70% of the observations for training and 30% for testing, bootstrapping the data with resampling, in line with the Data Perturbation Ensemble procedure [5]. Each computation returns 500 vectors of microRNAs sorted for decreasing “importance score”, i.e. the impact of each feature in the classification calculated by the algorithm used: for Gain Ratio the Entropy index was computed, for Random Forest we calculated a mean value between the *MeanDecreaseAccuracy* and the *MeanDecreaseGini* indices, and for the SVM-RFE the importance score was the mean accuracy returned from each classification of the model. In particular, the higher the importance score, the lower the “rank” that has been assigned to the feature. Then we applied an *aggregation* procedure to derive a “consensus” signature for every FS approach: we used a major voting-based aggregation strategy, in which the feature that at most occurred in a certain rank among all the 500 bootstraps, was finally assigned to that position in the “consensus” aggregated signature. For each panel of microRNAs, the top-200 scoring features have been maintained.

### 2.2 Stability Measures

The Kuncheva Index (KI), and the Percentage of Overlapping Gene/Features (POG) have been used to evaluate the consistency of the FS approaches used [5, 6]. In particular, we also used a pairwise measure of the KI, the S_tot_ statistic, as formulated by Abeel and colleagues [4] (Equation 1), in order to determine the stability between all the methods regardless of the aggregation strategy used [5]. These statistics were computed at incremental lengths of the signatures, starting from a minimum of 2 features to a maximum of 200, with an increase of 2 units for each recalculation.

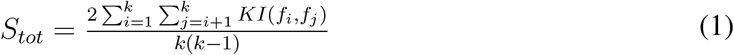

### 2.3 Differential Expression Analysis and DE-signatures

The DEA on the TCGA dataset, both for microRNA-Seq and RNA-Seq data, was performed starting from the raw counts, using the Exact Test, then DE features with a FDR *<*= 0.01 and a Log_2_FC cutoff set to |0.5| were maintained. To obtain a signature of DE-microRNAs, the Log_2_FC values were turned into their absolute value, and the microRNAs were sorted in decreasing abs(Log_2_FC) order (first 200 features maintained). The GEO validation dataset has been subjected to Limma [7] for DEA; the DE-signature of this dataset has been obtained using the same parameters and procedure applied for TCGA data.

### 2.4 Classification Performance analyses

In order to establish the ability of each signature to discriminate between healthy and cancer patients, a predictivity analysis was implemented in the pipeline on both the testing sub-dataset (TCGA) and validation dataset (GEO) for each of the four signatures, comprising FS panels and DE ones. We used a SVM classifier having a minimal configuration, with Radial Basis kernel “Gaussian”, C parameter set to 1, an optimal sigma chosen automatically by the function; we applied a 5-fold Cross-validation strategy on both tests. Finally, we computed the average values of Accuracy (ACC), K statistic (KK) and Matthews Correlation Coefficient (MCC) across the several folds and at several lengths of every signature as done in subsection 2.2.

### 2.5 SVM-RFE microRNA-signature Target detection

To identify the possible gene targets of the chosen microRNAs, we proceeded in four steps:

1. top-20 SVM-RFE microRNAs have been sorted in up and down-regulated in tumoral samples;
2. a DEA was performed on the RNA-Seq data to detect DE genes (FDR *<*= 0.05);
3. a Spearman correlation analysis between the expression of the microRNAs and that of the DE genes was applied to only maintain the up-genes that were inversely correlated with down-microRNAs and down-genes inversely correlated with up-microRNAs (rho *<*= -0.5);
4. all the experimental validated microRNAs’ gene targets have been collected and only the ones also DE-correlated have been maintained.

### 2.6 Network Centrality and hub genes identification

A correlation matrix (Spearman) of selected dysregulated genes has been used to build a graph gene Network: we retained only hub genes with a Kleinberg’s hub centrality score *>* 75 [8] and with a rho *>* 0.8 or rho *<* -0.6. An Over-representation analysis (ORA) 3 on the hub genes has been computed to explore the most enriched pathways from REACTOME database.4 FDR adjusted pValue threshold was set to 0.005.

## 3 Results

Each FS method applied returned 500 microRNAs signatures, sorted by decreasing importance score and then aggregated to finally have three “consensus” panels. Interestingly the top 3 microRNAs *hsa-mir-139, hsa-mir-96* and *hsa-mir-145* were maintained in all the panels, underlining the importance of these molecules in distinguishing tumoral from healthy samples.

### 3.1 The SVM-RFE is the most stable approach

From KI and POG computation on consensus panels, the SVM-RFE approach results to have the highest stability, in particular when the signature reaches a length of about 20 features. Even the S_tot_ index, computed on all the 500 signatures for each method before the aggregation step, shows the highest stability for the SVM-RFE (Figure 2)

**Figure 2:**
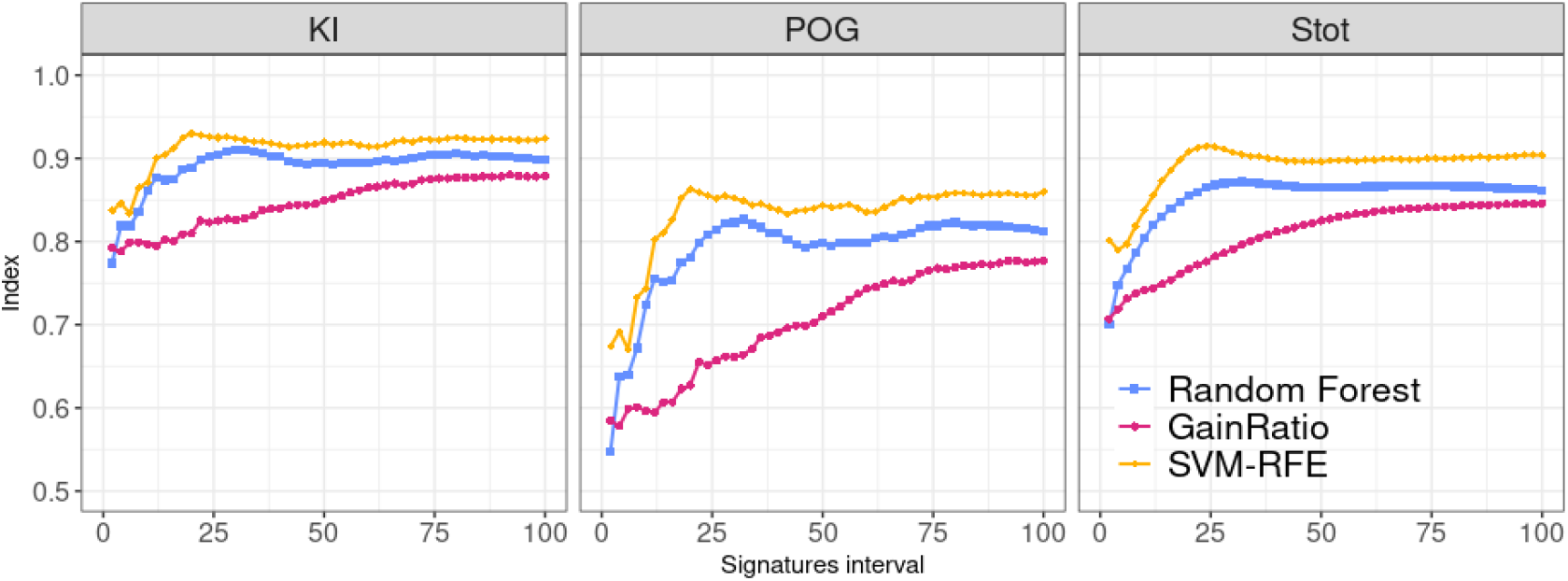
Comparison of Stability indices across the methods. KI: Kuncheva Index; POG: Percentage of overlapping Genes; S_*tot*_: pairwise KI measure.

### 3.2 The SVM-RFE signature performs better in classification than DE signatures

The classification performance analysis done on every single panel, returned for both the testing (TCGA) and the validation datasets (GEO), that the signature obtained with SVM-RFE has the highest predictive power. Figure 3 panel A, shows the average statistics across the 5 folds and for the top-20 microRNAs of each signature, since as described in subsection 3.1 this corresponds to the minimum length needed to maintain high stability: the SVM-RFE is clearly the best-performing signature. For the validation dataset, we selected the top-20 microRNAs of our signatures that were available and also in this case, the SVM-RFE returns a higher prediction ability. Investigating more deeply the first 20 microRNAs of the panels obtained, with particular attention to the most robust signature, we noticed some microRNAs shared across the methods but whose role in Breast Cancer is still not completely established: *hsa-mir-337, hsa-mir-378c* and *hsa-mir-483* (Figure 3 panel B).

**Figure 3:**
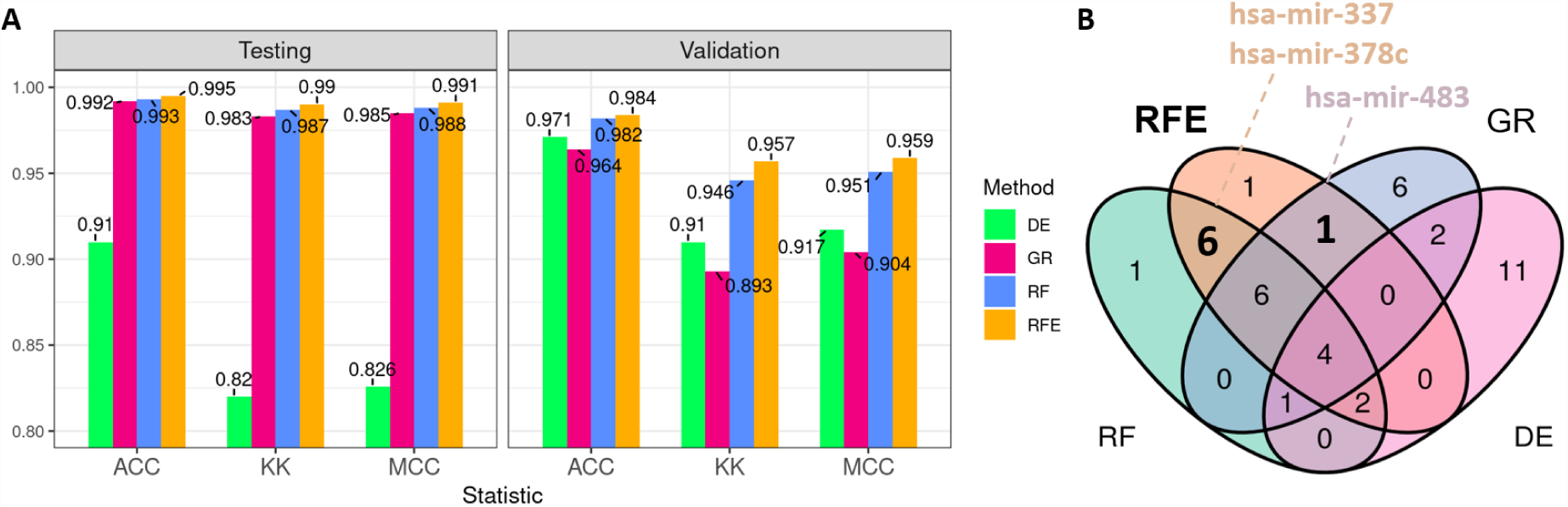
top-20-microRNAs classification performance and Venn graph: in (A) the barplot shows the mean statistics computed on testing sub-dataset and on external validation GEO dataset; in (B) a Venn graph of the top-20 microRNAs for each signature, where some interesting microRNAs of the top-20 of the SVM-RFE panel have been labeled. ACC: Accuracy; KK: K statistic; MCC: Matthews Correlation Coefficient; RFE: SVM-RFE; RF: Random Forest; GR: Gain Ratio; DE: Differential Expression

### 3.3 Network analysis uncovers putative key genes in the evolution of the disease

The SVM-RFE was therefore chosen for the construction of the network, then 40 up-regulated hub-genes (*CDC25, TPX2, KIF18B* and *CDCA3* representing the top 4) and 4 down-regulated hub-genes (*TGFBR2, CAV1, TNS1* and *LHFPL6*) have been maintained. The ORA performed on all the up-regulated targets enriches pathways related to the cell cycle and in particular to the transition to the M phase, as shown in Figure 4; in contrast, *TGFBR2* and *CAV1* are associated to *integrin-mediated cell adhesion* 5 and *TGF-****β*** *signaling pathway*6. By further investigating the role of these genes in Breast Cancer, we find that for some of them there is strong evidence of involvement in the disease: the study by Suo and coworkers highlights how *CDC25, TPX2* and *KIF18B* are strongly expressed in different cancer types as well as related to stemness in triple-negative patients [9]; furthermore, *TGFBR2* down-regulation is related to cancer progression as reported by Novitskiy and colleagues [10].

**Figure 4:**
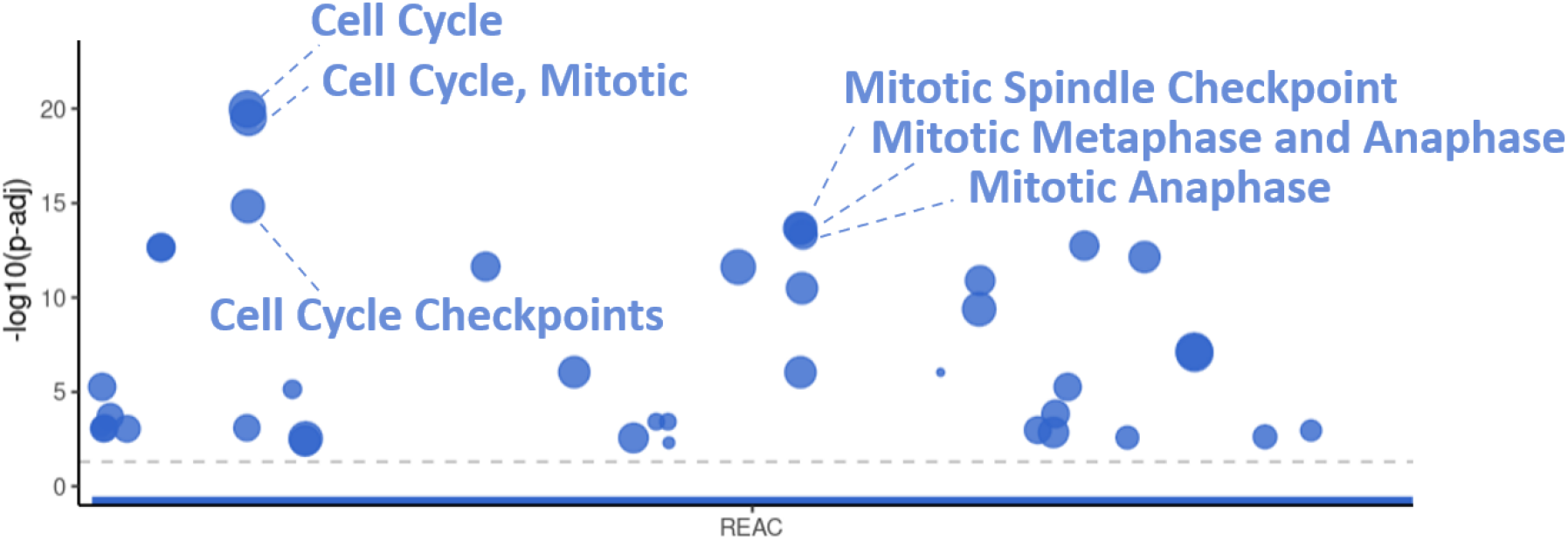
Bubble chart of Enrichment pathway analysis (ORA) of the 40 up-regulated hub-gene targets: the top 6 significant REACTOME pathways enriched have been labeled; pValue adjusted (FDR) threshold = 0.005 is indicated by the gray dotted line.

## 4 Conclusions

This study underlines as machine learning-based approaches represent a turning point in the field of biomarker research, indeed, SVM-RFE-derived consensus microRNAs signature got better performances in segregating tumoral from healthy patients when compared to DE-derived panels, both for testing and validation classification analyses. These results uncover microRNAs that can be underestimated by the DEA and which instead appear to have high discriminating power in both the datasets we analyzed. So we suggest a role for the top-20 SVM-RFE microRNAs as a panel of specific diagnostic biomarkers of Breast Cancer. Moreover, although our results are very preliminary and require further confirmation with *in wet* experiments, they assess the crucial importance of key microRNAs as *hsa-mir-139, hsa-mir-96* and *hsa-mir-145* and uncover new, putative biomarkers *hsa-mir-337, hsa-mir-378c* and *hsa-mir-483*, whose role in the disease is still not completely elucidated. Moreover, we characterized the contribution of some markers, such as *CDC25, TPX2* and *KIF18B*, in disease progression and evolution. For the down-regulated genes, it is crucial to better explore the meaning of their deactivation since these findings could lead to the discovery of key mechanisms which, if restored, could help downtrend the progression of the neoplasm.

## Conflict of interests

The authors declare that they have no conflict of interest.

## Footnotes

1 https://www.cancer.gov/ccg/research/genome-sequencing/tcga

2 https://www.ncbi.nlm.nih.gov/geo/

3 https://biit.cs.ut.ee/gprofiler/gost

4 https://reactome.org/

5 https://www.wikipathways.org/pathways/WP185.html

6 https://www.wikipathways.org/pathways/WP366.html

## Notes

### Competing Interest Statement

The authors have declared no competing interest.

### Summary of Updates

Image and text typos have been corrected

## References

[1] Y. Peng and C. M. Croce, “The role of MicroRNAs in human cancer,” Signal Transduction and Targeted Therapy, vol. 1, p. 15004, Jan. 2016.

[2] P. R. V. ml, and S. S., “Gain ratio based feature selection method for privacy preservation,” ICTACT Journal on Soft Computing, vol. 01, pp. 201–205, 04 2011.

[3] L. Breiman, “Random forests,” Machine Learning, vol. 45, no. 1, p. 5–32, oct 2001.

[4] T. Abeel, T. Helleputte, Y. Van de Peer, P. Dupont, and Y. Saeys, “Robust biomarker identification for cancer diagnosis with ensemble feature selection methods,” Bioinformatics, vol. 26, no. 3, pp. 392–398, 11 2009.

[5] Z. He and W. Yu, “Stable feature selection for biomarker discovery,” Computational Biology and Chemistry, vol. 34, no. 4, pp. 215–225, 2010.

[6] K. L. I., “A stability index for feature selection,” in Proceedings of the 25th Conference on Proceedings of the 25th IASTED International Multi-Conference: Artificial Intelligence and Applications. ACTA Press, 2007, p. 390–395.

[7] M. E. Ritchie, B. Phipson, D. Wu, Y. Hu, C. W. Law, W. Shi et al., “limma powers differential expression analyses for RNA-sequencing and microarray studies,” Nucleic Acids Research, vol. 43, no. 7, pp. e47–e47, Apr. 2015.

[8] J. M. Kleinberg, “Authoritative sources in a hyperlinked environment,” Journal of the ACM, vol. 46, no. 5, p. 604–632, sep 1999.

[9] H. D. Suo, Z. Tao, L. Zhang, Z. n. Jin, X. y. Li, W. Ma et al., “Coexpression network analysis of genes related to the characteristics of tumor stemness in triple-negative breast cancer,” BioMed Research International, vol. 2020, p. 7575862, 2020, publisher: Hindawi.

[10] S. V. Novitskiy, M. W. Pickup, A. E. Gorska, P. Owens, A. Chytil, M. Aakre et al., “TGF-β Receptor II Loss Promotes Mammary Carcinoma Progression by Th17-Dependent Mechanisms,” Cancer Discovery, vol. 1, no. 5, pp. 430–441, 10 2011.

